# Mitochondria decode firing frequency and coincidences of postsynaptic APs and EPSPs

**DOI:** 10.1101/2021.06.07.447340

**Authors:** Ohad Stoler, Alexandra Stavsky, Yana Khrapunsky, Israel Melamed, Grace Stutzmann, Daniel Gitler, Israel Sekler, Ilya Fleidervish

**Author notes:** co-corresponding authors Address for correspondence: Ilya A. Fleidervish, Department of Physiology and Cell Biology, Faculty of Health Sciences, POB 653, Ben Gurion University, Beer–Sheva 84105, Israel, Israel Sekler, Department of Physiology and Cell Biology, Faculty of Health Sciences, POB 653, Ben Gurion University, Beer–Sheva 84105, Israel. equal contribution (Ohad Stoler, Alexandra Stavsky).

## Abstract

Mitochondrial metabolism is critical for brain function. However, the mechanisms linking mitochondrial energy production to neuronal activity are elusive. Using whole-cell electrical recordings from Layer 5 pyramidal neurons in cortical slices and fluorescence imaging of cytosolic, mitochondrial Ca^2+^ indicators and endogenous NAD(P)H, we revealed ultra-fast, spike-evoked mitochondrial Ca^2+^ transients temporally similar to cytosolic Ca^2+^ elevations. We demonstrate that, whereas single or few spikes elicit the mitochondrial Ca^2+^ transients throughout the cell, their amplitude is differentially regulated in distinct neuronal compartments. Thus, these signals were prominent in the soma and apical dendrites and ∼3 times smaller in basal dendrites and axons. The spike firing frequency had a subtle effect on the amplitude of the cytosolic Ca^2+^ elevations but dramatically affected mitochondrial Ca^2+^ transients and NAD(P)H oxidation and recovery rates. Moreover, while subthreshold EPSPs alone caused no detectable Ca^2+^ elevation in dendritic mitochondria, the Hebbian coincidence of unitary EPSP and postsynaptic spike produced a localized, single mitochondrial Ca^2+^ elevation. These findings suggest that neuronal mitochondria are uniquely capable of decoding firing frequency and EPSP-to-spike time intervals for tuning the metabolic rate and triggering changes in synaptic efficacy.

---

Complex electrochemical and molecular processes underlying neuronal computation in the brain impose an immense, highly variable energy demand ^1,2^. For the neuronal circuit to function properly, the energy demand in all compartments of the individual neurons needs to be precisely matched by local, primarily mitochondrial ^3,4^, ATP production. During periods of enhanced neuronal activity, mitochondria accelerate ATP production via the feedforward mechanism by allowing the cytosolic Ca^2+^ elevations to propagate into the mitochondrial matrix ^2,5^. When cytosolic [Ca^2+^]_i_ rises, Ca^2+^ ions, powered by the steep mitochondrial membrane potential, flow into the mitochondrial matrix via the Ca^2+^ uniporter MCU ^6,7^, and are then extruded back into the cytosol by the mitochondrial Na^+^/Ca^2+^ antiporter NCLX ^8^. The mitochondrial Ca^2+^ elevation increases ATP production by activating at least three Krebs cycle enzymes ^9,10^. When Ca^2+^ signaling is disrupted, neuronal mitochondria fail to maintain the stable ATP concentration during enhanced activity periods ^5,11,12^. Moreover, dendritic mitochondria are critical for activity-dependent synaptic plasticity ^4^, but the link between the electrical activity, metabolism, and changes in synaptic strength is poorly understood.

Here, using whole-cell electrical recordings from Layer 5 pyramidal neurons in cortical slices and fluorescence imaging of cytosolic, mitochondrial Ca^2+^ indicators, we show that single or few spikes trigger rapidly rising and decaying mitochondrial Ca^2+^ elevations, with kinetics similar to cytosolic Ca^2+^ transients. Our evidence indicates that the mitochondria’s Ca^2+^ signaling and metabolic rate depend critically on spike firing frequency. We also show that, in dendrites, the coincidence of unitary EPSP and a backpropagating action potential produces a localized [Ca^2+^]_m_ transient, which, in addition to enhancing local ATP synthesis, could play a role in spike-time-dependent synaptic plasticity.

## Spike-elicited mitochondrial Ca^2+^ transients

To determine how neuronal electrical activity affects mitochondrial Ca^2+^ dynamics, we performed somatic whole-cell recordings from L5 pyramidal neurons expressing the mitochondria-targeted, neuron-specific Ca^2+^ indicator, mitoGCaMP6m. The intracellular solution was supplemented with the cytosolic Ca^2+^ indicator, Fura-2, that could be excited separately from mitoGCaMP6m. Series of thin, high-resolution optical sections over the vertical extent of the neuron, representing Fura-2 and mitoGCaMP6m fluorescence elicited by two-photon excitation at 760 and 960 nm, respectively, were used to reconstruct the morphology of the neuron and to reveal the position of the individual mitochondria within its soma and processes. To improve the temporal resolution of the optical signals, dynamic fluorescence measurements were obtained from the smaller regions of interest in soma, axon, and dendrites. In a typical experiment, two APs elicited by the injection of two brief current pulses via the patch pipette caused cytosolic and mitochondrial Ca^2+^ elevations in the soma of an L5 cell (**Figure 1a**). Whereas the cytosolic Ca^2+^ signals had relatively even intensity, the change in mitoGCaMP6m fluorescence occurred at “hotspots”, each representing an individual mitochondrion. The mitochondrial Ca^2+^ elevations began with a short delay after the beginning of the spike train, and they were observed only in the electrically active neurons. At the same time, the fluorescence of the nearby mitoGCaMP6m expressing mitochondria belonging to the non-active cells did not change (**Suppl. Figure 1**).

**Figure 1.**
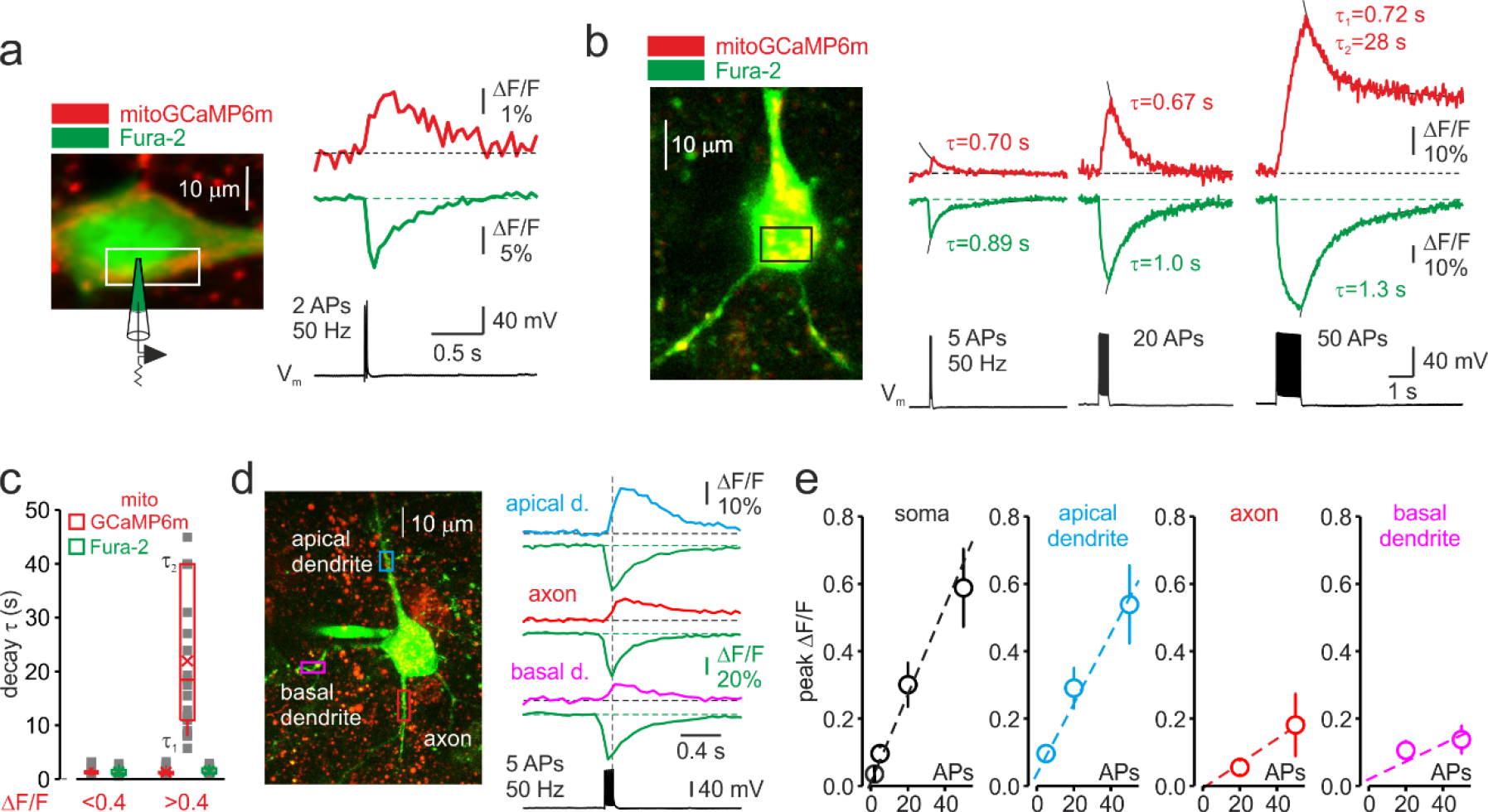
Trains of action potentials elicit mitochondrial Ca^2+^ elevations in soma and processes of L5 pyramidal neurons. **(a)** Changes in mitoGCaMP6m and Fura-2 fluorescence elicited by a train of two APs in a representative L5 pyramidal cell. *Left*, The image obtained by merging two optical sections through part of a L5 neuron at excitation wavelengths of 760 and 960 nm, eliciting the Fura-2 and mitoGCaMP6m fluorescence, respectively. *Right*, Somatic mitoGCaMP6m (red) and Fura-2 (green) ΔF/F transients elicited by a train of two APs. MitoGCaMP6m trace (red) is an ensemble average of 16 consecutive sweeps. **(b)** Small-amplitude mitoGCaMP6m transients decay rapidly. *Left*, Localization of the mitoGCaMP6m labeled mitochondria in a representative L5 neuron. The image was obtained by merging the fourteen, 1 µm thick optical sections at excitation wavelengths of 760 and 960 nm, eliciting the Fura-2 and mitoGCaMP6m fluorescence, respectively. The yellow color indicates colocalization of the Fura-2 (green) and mitoGCaMP6m (red) fluorescence, revealing the position of the mitochondria. The rectangle indicates the region of interest from which the optical sweeps were obtained. *Right*, somatic mitoGCaMP6m (red) and Fura-2 (green) transients elicited by 5, 20, and 50 APs in a cell shown on the left. Black lines are the best single or double exponential fits of the decay. Notice that after the relatively small [Ca^2+^]m elevations evoked by 5 or 20 APs, the Ca^2+^ clearance is rapid and follows the single exponential time course (*τ* ∼0.7 s). The decay of larger transients evoked by 50 APs is biexponential (*τ*1 of ∼0.7 s and *τ*2 of ∼28 s). **(c)** The decay time course of the small-amplitude (peak ΔF/F<0.4) and large-amplitude (peak ΔF/F>0.4) mitoGCaMP6m transients. Smaller amplitude transients (n=25) decayed mono-exponentially, whereas the decay of the larger transients (n=22) followed the bi-exponential time course. Each dot (grey) represents a decay time constant (*τ* for monoexponential decays, *τ* 1 and *τ* 2 for bi-exponential decays) of transients elicited by 5-50 APs in 42 neurons. Box plots represent the 25–75% interquartile range, and the whiskers expand to the 5–95% range. A horizontal line inside the box represents the median of the distribution, and the mean is represented by a cross symbol (X). **(d)** The amplitude of spike-evoked mitoGCaMP6m fluorescence transients varies between different neuronal processes. *Left*, Localization of the mitoGCaMP6m labeled mitochondria in a representative L5 pyramidal neuron. The rectangles indicate the regions within the apical dendrite (cyan), basal dendrite (magenta), and axon initial segment (red) from which fluorescence measurements were obtained. *Right*, Mitochondrial and cytosolic (green) Ca^2+^ transients elicited by a train of 5 APs at 50 Hz in different neuronal compartments. **(e)** Mean peak ΔF/F of the mitoGCaMP6m transients as a function of the number of action potentials in soma, apical and basal dendrites, and axon initial segment. Shown are mean values ± SE (n=5-29).

The AP-elicited mitochondrial Ca^2+^ elevations required Ca^2+^ influx via the voltage-gated Ca^2+^ channels, as bath application of the Ca^2+^ channel blocker, Cd^2+^ (200 µM) inhibited both cytosolic and mitochondrial Ca^2+^ transients (**Suppl. Figure 2**). Previous studies have shown that Ca^2+^ depletion of endoplasmic reticulum (ER) in cultured neurons does not influence mitochondrial Ca^2+^ elevations ^5^. Consistent with these results, we found that blockade of ER Ca^2+^ channels by Ryanodine (100 µM) and Dantrolene (100 µM) (**Suppl. Figure 3**) had no significant effect on mitochondrial Ca^2+^ transients. The mitochondrial Ca^2+^ channel, MCU, is a primary route for Ca^2+^ entry to the mitochondrial matrix ^6,7,9^. In our experiments, blockade of the MCU by intracellular application of Ruthenium red almost completely abolished the mitochondrial Ca^2+^ signals (**Suppl. Figure 4**) whereas little to no change in amplitude of cytosolic Ca^2+^ elevations was detected.

An increase in the number of spikes produced progressively larger mitoGCaMP6m transients in the soma (**Figure 1b**) and the apical dendrite (**Suppl. Figure 5**) with a progressively slower rising phase. Thus, the mito-Ca^2+^ signals elicited by 50 APs grew throughout the spike train duration, reaching a peak ΔF/F value of 59 ± 11% (n=29) and 54 ± 11% (n=22) for soma and apical dendrite, respectively, at 120 ± 18 ms (n=10) after the train end. In contrast with previous reports ^5,13^, mito-Ca^2+^ transients elicited by 2-20 spikes decayed as rapidly or even faster than cytosolic Ca^2+^ transients. However, following large elevations, [Ca^2+^]_m_ remained high for tens of seconds, reaching the resting level long after the cytosolic Ca^2+^ concentration completely recovered. We systematically examined the relationship between the peak amplitude and the decay rate of the mito-Ca^2+^ transients in 42 pyramidal neurons (**Figure 1c, Suppl. Figure 6**). Decay of smaller transients (peak ΔF/F <40%) was always monoexponential, with τ=0.86±0.11 s (n=25) as well as decay of some middle-sized (ΔF/F 40-75%) transients (τ=0.82±0.10 s, n=10). The decay of other middle size and large (ΔF/F >75%) transients was double exponential, with τ1=0.86±0.13 s and τ2=22.0±2.85 s (n=23). The decay of cytosolic Ca^2+^ transients always followed a single exponential time course which was characterized by τ=1.03±0.14 s (n=22) for small and τ=1.19±0.4 s (n=20, p=0.43) for large transients.

To elucidate the differences in mitochondrial Ca^2+^ signaling between distinct neuronal compartments, we monitored mitoGCaMP6m fluorescence in ∼10 µm long regions of interest in basal, apical dendrites, and axon initial segments (AISs) during trains of five APs (**Figure 1d**). While the amplitude and time course of the cytosolic Ca^2+^ responses in all these compartments was similar, the magnitude of mitochondrial signals was remarkably polar. The amplitude of the mitochondrial Ca^2+^ elevations in apical dendrite was as prominent as in the soma. In contrast, in the AIS and thin basal dendrites, the mito-Ca^2+^ responses were dramatically smaller.

We next sought to evaluate the relationship between the number of APs and the mean peak amplitude of the mitoGCaMP6m transients in the soma, AIS, apical and basal dendrites of 29 neurons (**Figure 1e**). The steepness of this relationship was greater in soma and apical dendrites (1.3% and 1.1% ΔF/F per spike, respectively) compared with AIS and basal dendrites (0.4% and 0.3% ΔF/F per spike, respectively). The compartmental differences in mitochondrial signals were not due to a different magnitude of cytosolic Ca^2+^ elevations. The pattern of [Ca^2+^]_i_ during the neuronal activity is known to be complex. However, the differences in cytosolic Ca^2+^ levels were subtle and they poorly correlated with the magnitude of mitochondrial Ca^2+^ elevations. For example, the peak ΔF/F amplitude of Fura-2 transients elicited by twenty spikes was 26±3.6% (n=22) for soma, 37±4% (n=13) for apical dendrite, 38±4% (n=13) for basal dendrites, and 25±4% in the AIS (n=7). The compartmental differences in magnitude of the mitoGCaMP6m transients could, at least partially, be explained by the lower expression level of fluorescence probe in the mitochondria localized within the thinner neuronal processes. This seems to be unlikely, however. Thus, in all neuronal compartments, maximal mitoGCaMP6m fluorescence of the individual mitochondria following prolonged depolarization of the neuronal membrane was similar (**Suppl. Figure 7**). Our evidence, therefore, points to the existence of an as-yet-unidentified mechanism that differentially regulates the mitochondrial Ca^2+^ entry in distinct neuronal compartments.

## Frequency-dependent amplification of mitochondrial Ca^2+^ elevations

Next, we examined whether mitochondrial Ca^2+^ elevations are sensitive to firing frequency. **Figure 2a** shows an optical recording from the soma of a representative L5 neuron in which cytosolic and mitochondrial Ca^2+^ transients were elicited by trains of twenty APs at 20, 50, and 100 Hz. As spike frequency increased, the rise of cytosolic Ca^2+^ transients became progressively faster and their peak amplitude modestly increased (**Suppl. Figure 8)**. In contrast, the mitochondrial Ca^2+^ signals showed a very different frequency dependence. Firing at a frequency of 20 Hz or lower elicited a minimal elevation in [Ca^2+^]_m_, whereas the response to spikes at a frequency of 50 Hz or higher was dramatically greater. The steep frequency dependence of the mitochondrial signals was observed in all neuronal compartments, including somas, apical and basal dendrites, making it unlikely that it reflects the frequency-dependent failure of AP backpropagation ^14^. The frequency-dependent amplification of mitochondrial Ca^2+^ transients was observed in all 21 neurons tested with either 50, 20 or 5 APs (**Figure 2b**). The mean ratio of peak amplitudes of the mitochondrial Ca^2+^ transients elicited by trains of spikes at 50 and 20 Hz was larger when neurons were subjected to longer (3.36±0.46 times, n=22 for 50 APs) than to shorter (1.74±0.18 times, n=5 for 5 APs) trains (**Figure 2c**). Systematic varying of AP frequency in a range from 10 to 100 Hz revealed that the peak amplitude of mito-Ca^2+^ transients behaved as Bolzmannian function of the frequency, with mean half-amplitude of 38 Hz and steepness of 22 Hz^-1^ (n=12).

**Figure 2.**
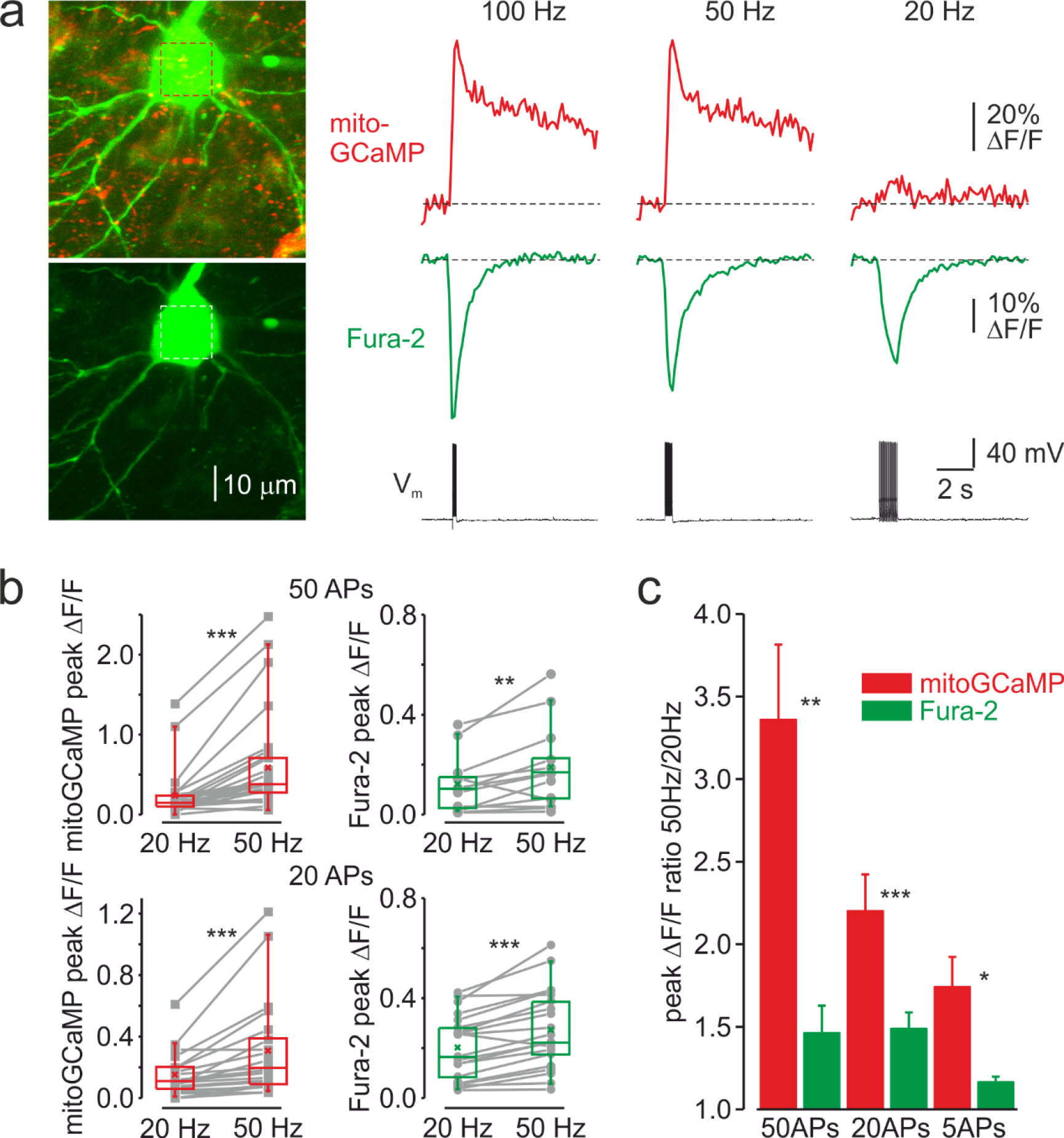
Frequency-dependent amplification of the mitochondrial Ca^2+^ elevations. **(a)** Cytosolic and mitochondrial Ca^2+^ elevations elicited at the soma of a representative pyramidal neuron by trains of 20 APs at 100, 50, and 20 Hz. *Left*, The top image is mitoGCaMP6m labeled mitochondrial map of a representative L5 pyramidal neuron, obtained by merging the Fura-2 and mitoGCaMP6m fluorescence (for detail, see Figure 1). The bottom image is the maximum intensity Z-projection representing the Fura-2 fluorescence only. *Right*, The mitoGCaMP6m (red) and Fura-2 (green) ΔF/F transients elicited at the soma by the spike trains at indicated frequency. Notice that, at 20 Hz, the train of spikes produced almost no mitochondrial Ca^2+^ elevation, while the amplitude of the cytosolic Ca^2+^ transient changed little as a function of spike frequency. **(b)** Peak amplitude of mitoGCaMP6m (red) and Fura-2 (green) ΔF/F transients elicited by a train of 50 (top) or 20 (bottom) APs at a frequency of 20 and 50 Hz. The grey lines connect the paired values obtained from the same individual neuron at two firing frequencies. Box plots represent the 25–75% interquartile range, and the whiskers expand to the 5–95% range. A horizontal line inside the box represents the median of the distribution, and the mean is represented by a cross symbol (X). **(c)** Mean ratio of peak amplitudes of mitoGCaMP6m (red) and Fura-2 (green) ΔF/F transients elicited by trains of 50, 20, and 5 APs at 50 and 20 Hz frequency.

## Frequency-dependent acceleration of the mitochondrial NAD(P)H metabolism

To determine whether the frequency-dependent amplification of Ca^2+^ signals in the mitochondria affects their metabolic activity, we monitored the changes in NAD(P)H autofluorescence elicited by trains of brief, just suprathreshold antidromic stimuli at a different frequency (**Figure 3a**). Whole-cell recording from a single pyramidal neuron within the region of interest was obtained to tune the stimuli intensity such that each stimulus would elicit only one AP. In cortical neurons, NAD(P)H signals primarily reflect changes in mitochondrial NAD(P)H pool ^2^. In response to electrical stimulation, we observed a negative deflection in the NAD(P)H autofluorescence (“dip”) which indicates an increased rate of electron transfer reflected in NAD(P)H oxidation, followed by a positive transient (“overshoot”) which indicates Krebs cycle dependent replenishment of NAD(P)H pool. At higher stimulation frequency, the magnitude of both dip and overshoot of the NAD(P)H signals were enhanced, consistent with the previous reports that the rates of NAD(P)H oxidation and synthesis are dependent on the Ca^2+^level in the mitochondrial matrix ^2^. Spatio-temporal analysis of the NAD(P)H autofluorescence dynamics at two stimulation frequencies (**Figure 3b**) revealed that the changes in the fluorescence were spatially restricted to Layer 5 of a single cortical column and that the frequency-dependent amplification of both dip and overshoot amplitude was relatively uniform within the stimulated region. A comparison of NAD(P)H autofluorescence changes in response to a train of 50 stimuli at 20 Hz and 50 Hz in ten cortical slices (**Figure 3c**) revealed the same frequency dependence as with [Ca^2+^]_m_. The dip amplitude increased from -15±2 a.u. at 20 Hz to -21±2 a.u. at 50 Hz (n=15 ROIs, p<0.001), the overshoot’s peak amplitude increased from 8±2 a.u. to 19±3 a.u. (p<0.001) and the overshoot area increased from 156±46 a.u.·s to 441±71 a.u.·s (p<0.001), respectively. Because the glial responses might partially contaminate NAD(P)H signals obtained in the cortical slices, we tested the frequency dependence of NAD(P)H responses in the in Stratum pyramidale of CA1 area of the hippocampus (**Suppl. Figure 9**) which predominantly contains neuronal cell bodies. As in the neocortex, NAD(P)H signals in response to trains of stimuli delivered to the Stratum oriens were significantly enhanced at the higher stimulation frequency.

**Figure 3.**
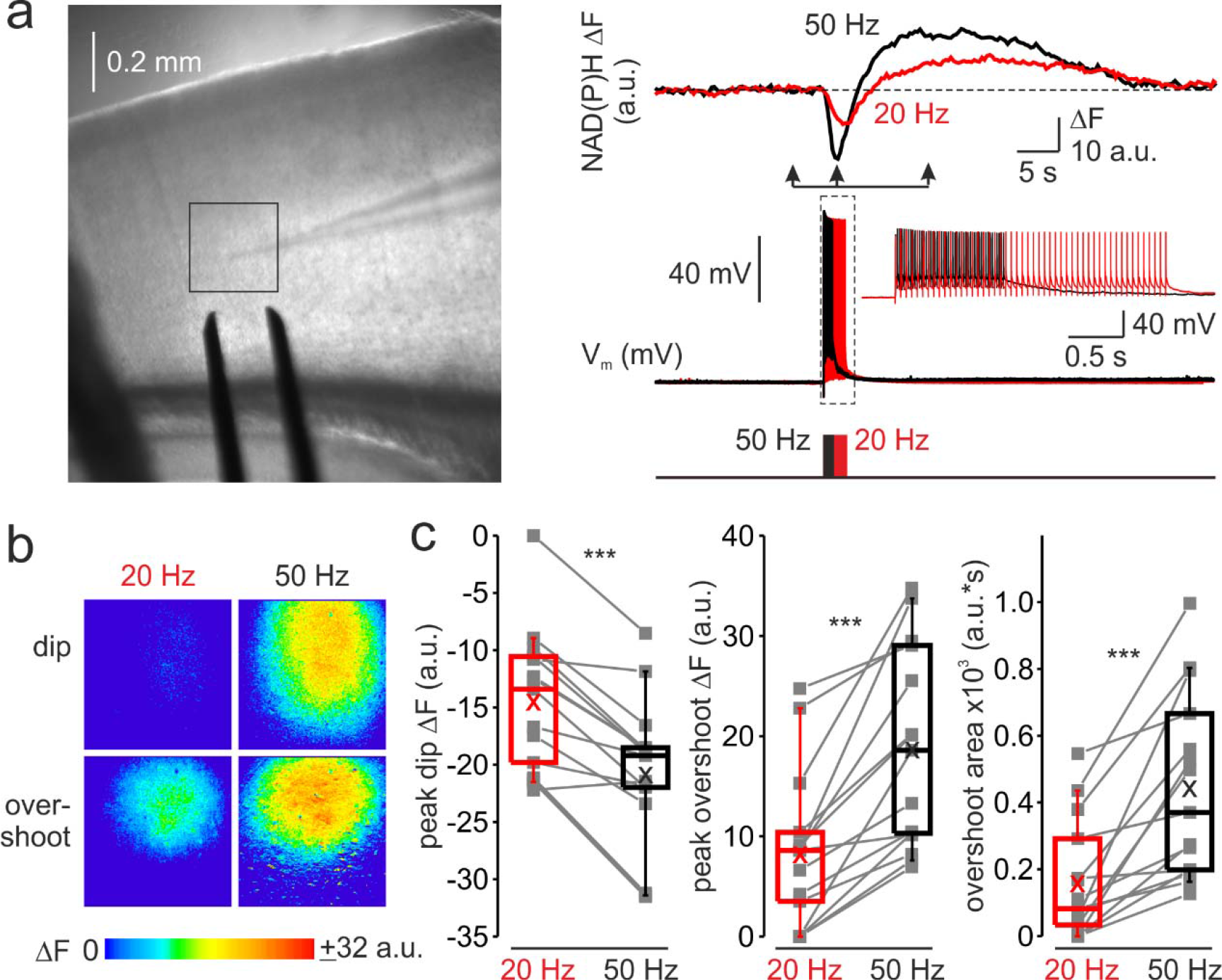
Frequency-dependent amplification of spike elicited changes in mitochondrial NAD(P)H auto-fluorescence. **(a)** In a representative cortical slice, changes in NAD(P)H fluorescence in response to extracellular stimuli trains depend on stimulation frequency. *Left*, DIC image of a coronal slice during the electrical and optical recording. The rectangle indicates the region from which the auto-fluorescence measurements were obtained. The stimuli were delivered via the bipolar electrode placed on the white-grey matter border, and the whole-cell recording was obtained from an L5 neuron within the same cortical column. *Right*, The membrane potential and optical traces evoked by trains of 50 just suprathreshold stimuli at 50 Hz (black) and 20 Hz (red). Notice that both dip and overshoot of the NAD(P)H signals are more prominent at 50 Hz. *Inset*: Stimuli intensity was carefully adjusted to elicit only a single AP per stimulus. **(b)** The amplitude of NAD(P)H signals depends on stimulation frequency, whereas their spatial extent does not. Shown are pseudocolor maps of change in the NAD(P)H fluorescence between the times marked by the arrowheads in **a**. **(c)** Higher frequency stimulation causes an increase in the magnitude of the dip and of the overshoot of the NAD(P)H signal. Box plots representing the peaks of the dip (left), the peaks of the overshoot (middle), and the area of the overshoot (right) of the NAD(P)H signals evoked by trains of 20 Hz (red) and 50 Hz (black) stimuli. The grey lines connect the paired values obtained from the same cortical regions at two firing frequencies (n=15 ROIs, ten cortical slices, three mice). Box plots represent the 25–75% interquartile range, and the whiskers expand to the 5–95% range. A horizontal line inside the box represents the median of the distribution, and the mean is represented by a cross symbol (X).

## Localized dendritic [Ca^2+^]_m_ elevations elicited by the coincidence of postsynaptic AP and EPSP

We next sought to elucidate how synaptic activity affects mitochondrial Ca^2+^ dynamics. After filling the cell for ∼20 min to allow the diffusion of Fura-2 into the dendrites, we positioned the bipolar electrode close to an apical or basal dendrite. Delivery of a single brief stimulus (0.1 ms), with its amplitude adjusted to keep the subsequent EPSP below the threshold for postsynaptic spike generation produced no detectable cytosolic or mitochondrial Ca^2+^ signals (n=7). The single AP elicited by brief somatic current pulse injection, however, produced a cytosolic but no mitochondrial Ca^2+^ response in the dendrites. Remarkably, the coincidence of the EPSP and backpropagating AP synergized to elicit a robust cytosolic and mitochondrial Ca^2+^ response (**Figure 4a**). While most dendritic mitochondria were silent, the EPSP and AP coincidence created single localized mitochondrial Ca^2+^ “hotspots” with peak ΔF/F amplitude of 6.4 ± 0.8% (n=8) in the dendritic regions of interest. We interpreted the appearance of these hotspots as evidence for a highly restricted, probably single mitochondrial Ca^2+^ elevation, in the vicinity of the active spine. The unitary character of the mitochondrial signals under this experimental paradigm can be explained by a spatial sparseness of the synapses formed by the presynaptic fiber on the dendrites of the postsynaptic cortical cell ^15^ such that only one out of a few currently active spines could be found in the relatively short segment of a dendritic branch that we examined. The cytosolic Ca^2+^ elevation, measured in the same hotspot at which the mitoGCaMP6m signal was detected, was not significantly larger than in the nearby dendrite. The failure to see the cytosolic Ca^2+^ “hotspots” is, most probably, due to temporal and amplitude resolution of our optical recording that was insufficient to reveal Ca^2+^ elevation in the tiny volume between the spine neck and the mitochondrion during the fast single spine Ca^2+^ transient ^16,17^. We extended our analysis by measuring the cytosolic and mitochondrial Ca^2+^ signals elicited by 20 unpaired APs and APs paired with EPSP. In both apical and basal dendrites, the paired APs produced a significantly larger mitochondrial signal. In contrast, the cytosolic Ca^2+^ elevation amplitude was not significantly different for unpaired and paired stimulation (**Figure 4b** and **c**). Hence, the peak amplitude of the mitoGCaMP6m transients elicited by paired APs was 2.73±0.32 (n=8) and 2.11±0.32 (n=9) times higher than of those evoked by the unpaired APs in the basal and apical dendrites, respectively (**Figure 4d**). The ability of postsynaptic neurons to generate an AP was crucial for triggering the dendritic mito-Ca^2+^ transients. Thus, intracellular dialysis with a solution containing a blocker of voltage-gated Na^+^ channels, QX-314, which prevented the firing of the postsynaptic cells ^18^ while producing only a minor effect on the EPSP generation, dramatically reduced the amplitude of synaptically evoked mito-Ca^2+^ transients (**Suppl. Figure 10a, b**). It is well established that the main route of Ca^2+^ entry into the spines is via the NMDARs ^17,19^, which require glutamate and depolarization to relieve the Mg^2+^ blockade ^20^. Blockade of NMDARs by APV (50 µM) almost completely and reversibly abolished the mitoGCaMP6m transients evoked by the paired APs in the dendrites (**Suppl. Figure 10c, d**).

**Figure 4.**
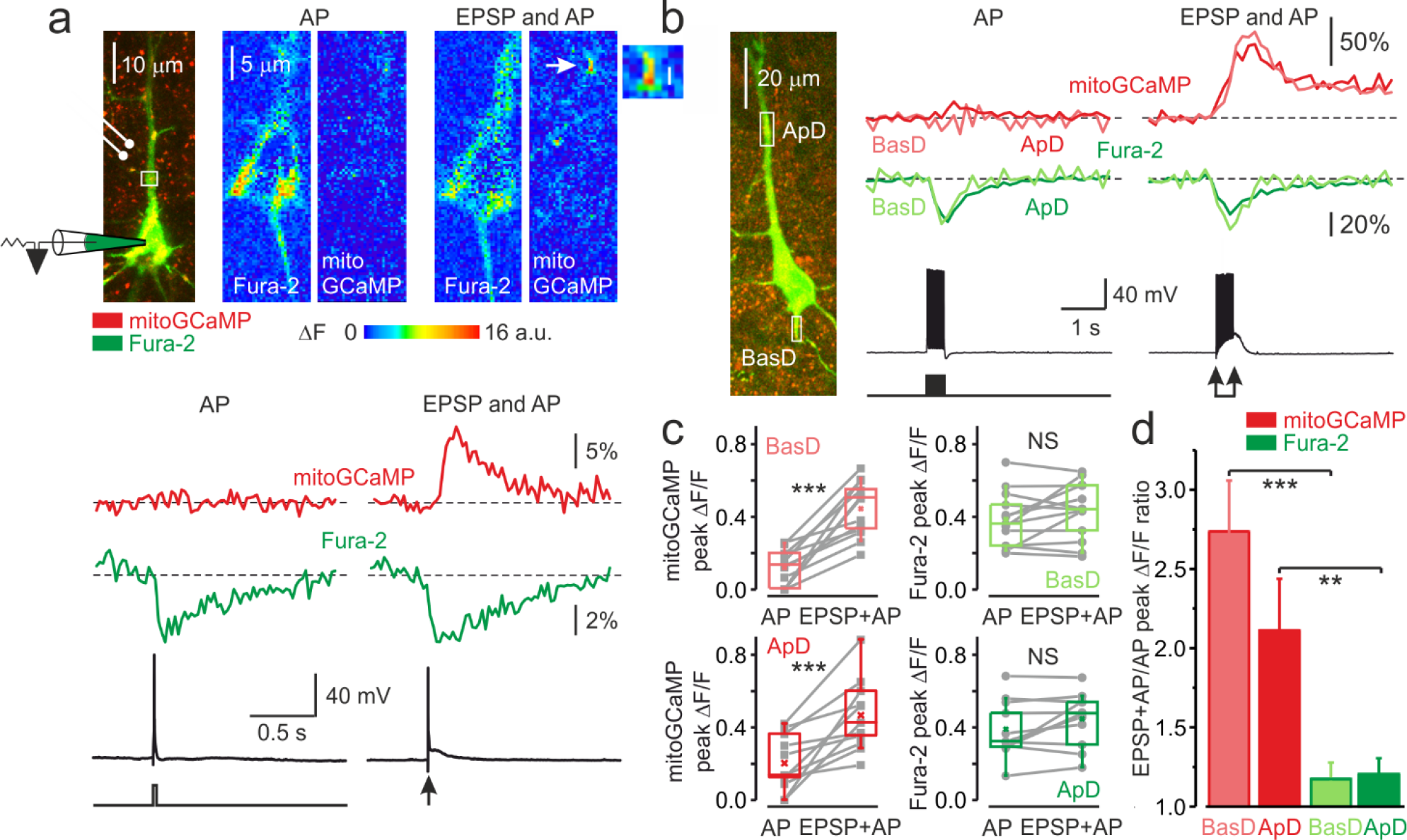
The coincidence of postsynaptic action potential and EPSP induces localized mitochondrial Ca^2+^ elevations in the dendrites. **(a)** Action potential elicited by a single, just suprathreshold synaptic stimulus induces a large, spatially restricted increase in dendritic mitoGCaMP6m fluorescence, whereas action potential evoked by a 5-ms current pulse (600 pA, cell body injection) had no such effect. *Top, Left*: Mitochondrial map in a representative L5 pyramidal neuron, obtained by merging the Fura-2 and mitoGCaMP6m fluorescence (for detail, see Figure 1). The rectangle indicates the region within the apical dendrite from which fluorescence measurements were obtained. *Right:* Pseudocolor maps of change in the mitoGCaMP6m and Fura-2 fluorescence in response to a synaptically and current pulse evoked AP. *Bottom*, The mitoGCaMP6m (red) and Fura-2 (green) ΔF/F transients measured from regions of interest as indicated in the upper panel and the somatic membrane potential trace. Arrow indicates a hotspot at which the synaptic stimulus elicited a mitochondrial Ca^2+^ transient. The transients are ensemble averages of 50 sweeps. **(b)** Action potentials elicited by a train of synaptic stimuli produce larger mitoGCaMP6m signals in the dendrites than APs elicited by the injection of a train of current pulses. *Left*: Mitochondria map in a representative L5 pyramidal neuron, obtained by merging the Fura-2 and mitoGCaMP6m fluorescence. The rectangles indicate the regions within the apical and basal dendrites from which fluorescence measurements were obtained. *Right*, Comparison of ΔF/F mitoGCaMP6m transients elicited in the apical (red) and basal dendrite (rose) by twenty suprathreshold synaptic stimuli at 50 Hz and by twenty brief current pulses delivered to the soma. The green and light green traces are ΔF/F Fura-2 transients in apical and basal dendrites, respectively. The black traces represent somatic membrane potential. **(c)** Peak amplitude of mitoGCaMP6m and Fura-2 ΔF/F transients elicited in basal (top) and apical dendrites (bottom) by a train of twenty suprathreshold current pulses or synaptic stimuli at a frequency of 50 Hz. The line connects the paired values obtained from the same individual neuron at two stimulation modalities. Box plots represent the 25–75% interquartile range, and the whiskers expand to the 5–95% range. A horizontal line inside the box represents the median of the distribution, and the mean is represented by a cross symbol (X). **(d)** Mean ratio of peak amplitudes of mitoGCaMP6m (rose, red) and Fura-2 (light green, green) ΔF/F transients elicited in dendrites by suprathreshold synaptic stimuli (EPSP+AP) and current pulses (AP). Data obtained from 8-13 individual neurons.

Using simultaneous electrical recordings, cytosolic and mitochondrial Ca^2+^ imaging in L5 pyramidal neurons, we found that relatively rare, singular spike firing, the firing mode associated with cortical circuit-based information processing in vivo ^21^, produces fast-rising, rapidly decaying mitochondrial Ca^2+^ transients in all neuronal compartments. These Ca^2+^ elevations, dramatically more rapid than those previously described in central neurons ^5,13^, were “tightly” coupled to electrical activity and cytosolic Ca^2+^ transients. In contrast to cytosolic Ca^2+^ signals, the summation of the unitary mitochondrial Ca^2+^ transients was highly non-linear. Thus, the mitochondria are capable of faithfully decoding the neuronal firing frequency into Ca^2+^ signals, which trigger changes in metabolic activity. Our findings about the prolonged clearance of Ca^2+^ ions from the mitochondrial matrix following the long periods of robust firing are consistent with previous observations ^5^. However, such intense firing is not typically observed in cortical pyramidal cells under physiological conditions, although it may occur during various neurological diseases ^22,23^ contributing to their progression. A recent study suggests that the metabolic activity of mitochondria plays a pivotal role in synaptic plasticity ^4^. How neuronal activity is linked to this process is poorly understood, however. Our results suggest that mitochondria can detect the Hebbian time coincidences between the pre- and postsynaptic spikes ^24^. The resultant Ca^2+^ elevations in the mitochondrial matrix could be an essential part of the cascade of events underlying spike-time-dependent synaptic plasticity. Our data indicate that the unique ability of the mitochondria to decode firing frequency and Hebbian timing code of neuronal activity make this organelle a long-thought link between the firing pattern, metabolism, and plasticity.

## Materials and Methods

### Experimental animals

All experiments were approved by the Animal Care and Use Committee of Ben Gurion University of the Negev. C57BL6 mice were obtained from Envigo (Israel).

### Viral constructs production and purification

cDNA of 2MT-GCaMP6m (mitoGCaMP6m) was subcloned by restriction/ligation (restriction enzymes and T4-ligase were from Fermentas/Thermo Scientific Life Science Research) into a plasmid containing adeno-associated virus 2 (AAV2) inverted terminal repeats flanking a cassette consisting of the neuronal-specific human synapsin 1 promoter (hSyn), the woodchuck post-transcriptional regulatory element (WPRE) and the bovine growth hormone polyA signal.

Viral particles were produced in HEK293T cells (ATCC) as previously described ^25^, using pAdDelta5 helper plasmids (a kind gift from Dr. Adi Mizrahi) and the pAAV2/9n plasmid (Addgene #112865) which encode the rep/cap proteins of AAV2 and AAV9, respectively. Viral particles were then purified over iodixanol (Sigma-Aldrich) step gradients and concentrated using Amicon filters (EMD). Virus titers were measured by determining the number of DNase I–resistant vector genomes (vg) using qPCR with a linearized genome plasmid as a standard ^26^.

### Stereotaxic injections

Mice at the age of P21-25 were deeply anesthetized with Ketamine/Xylazine and then stereotactic bilateral injections were performed into the Layer 5 of the somatosensory cortex using a microliter syringe (Hamilton, Israel) at a rate of 0.25 μl/minute, with 500 nl of AAV9-hSyn-Mito-GCaMP6m containing 1×10^10^ vg. After the injection, the needle was left in place for additional 3 minutes before being slowly removed from the brain. Coordinates for injections were (in mm): 4.1 rostral to lambda, ± 1.8 left/right of midline, -0.5 ventral to the pial surface.

### Acute coronal brain slices preparation

Coronal slices were prepared from mice three weeks post-injection (at the age of 6 to 7 weeks). The 300 µm thick coronal cortical or horizontal hippocampal slices were prepared using standard techniques, as previously described ^27,28^. Mice were anesthetized with isoflurane (5%) and decapitated. The slices were cut on a vibratome (VT1200; Leica) and placed in a holding chamber containing oxygenated artificial cerebrospinal fluid (ACSF) at room temperature; they were transferred to a recording chamber after more than 1 h of incubation. The composition of the ACSF (in mM): 124 NaCl, 3 KCl, 2 CaCl_2_, 2 MgSO_4_, 1.25 NaH_2_PO_4_, 26 NaHCO_3_, and 10 glucose; pH 7.4 when saturated with 95% O_2_/CO_2_.

### Fluorescence imaging

Most experiments were performed on L5 pyramidal neurons in somatosensory neocortical slices. The cells were viewed with a 40 or 60 × Olympus water-immersion lens of Ultima IV two-photon microscope (Bruker) equipped with a Mai Tai Deep See pulsed laser (Spectra-Physics). MitoGCaMP6m was excited at 940-950 nm. L5 pyramidal cells with low resting fluorescence that responded to electrical stimulation delivered via the nearby placed bipolar electrode were selected for whole cell recording (see below). In order to measure the cytosolic Ca^2+^ transients, the intracellular solution was supplemented by Ca^2+^ indicator, Fura-2 (100 µmol/l). The indicator was selected to minimize the interference with the mitoGCaMP6m fluorescence measurements. The Fura-2 fluorescence was elicited by two-photon excitation at 780 nm, and it declined as a function of the cytosolic Ca^2+^ concentration. The neuronal morphology and the labelled mitochondria localization was obtained by scanning a Z-series of 30-40 high resolution images at interval of 0.5 µm. The dynamic mitoGCaMP6m and Fura-2 imaging were performed from small regions of interest at frame rate of 10-50 Hz.

### Electrophysiology

Somatic whole-cell recordings were obtained using patch pipettes pulled from thick-walled borosilicate glass capillaries (1.5-mm outer diameter; Science Products, Germany). All recordings were at 30±0.5°C maintained with a temperature control unit (Luigs & Neumann, Rattingen). For current-clamp experiments the pipette solution contained (in mM): 130 K–gluconate, 6 KCl, 2 MgCl_2_, 4 NaCl, and 10 Hepes, with pH adjusted to 7.25 with KOH. Pipettes had resistances of 5–7 MΩ when filled with this solution supplemented with Fura-2 (Molecular Probes). Recordings were made using a Multiclamp 700B amplifier (Molecular Devices) equipped with CV-7B headstage (Molecular Devices). Data were low–pass–filtered at 10 kHz (−3 dB, single-pole Bessel filter and digitized at 20 kHz using Digidata 1322A digitizer driven by PClamp 10 software (Molecular Devices). Care was taken to maintain the access resistance below 10 MΩ.

### Wide-field fluorescence imaging

The NAD(P)H auto-fluorescence signals were obtained using a 40× water-immersion lens (Olympus) in a BX51WI microscope (Olympus). The fluorescence was excited by using a high-intensity LED device (385 ± 4 nm, Prizmatix), and the emission was collected by using a modified Olympus U-MNU2 filter set (DC=400 nm; EM=420 nm). Images were collected with the Orca-flash4.0 CMOS camera (Hamamatsu), using a pixel binning of 512×512, at a rate of 300 ms per frame. A bipolar stimulating electrode (WPI, 0.01 MΩ) was placed ∼100 µM below the region of interest, at the L-5/6 border in cortical slices or in CA1 stratum oriens in the hippocampal slices. The 0.1 ms long extracellular stimuli were delivered using an optically coupled stimulus isolation unit (A.M.P.I) driven via the pClamp 10 software. Somatic whole-cell recordings (see above) were made from a pyramidal neuron in the middle of the region of interest. The stimulation intensity was carefully controlled so that each stimulus triggered only a single antidromic spike with a latency of <1 ms post-stimulus. The baseline fluorescence was kept around 1500 a.u. throughout the experiments by regulating the intensity of LED emitted light.

## Data analysis

Electrophysiological data analysis was accomplished using pCLAMP10 software (Molecular Devices) and Origin 6.0 (OriginLab). The figures were created using CorelDraw X7 suite (Corel Corporation).

## Statistical analysis

If not otherwise noted, data are expressed as mean ± SE. Student *t*-test for paired or unpaired data was used for statistical analysis.

## Acknowledgements

This research was supported by The Israel Science Foundation (grant No. 1384/19 for IF and grant No. 1310/19 for DG), a US National Institutes of Health grant (RF1 AG065628).

## Author contributions

OS performed the electrical and imaging experiments, AS produced the virus and performed stereotactic injections, YK, IM, GS and DG performed data analysis, IS and IF designed the study and wrote the paper.

## Competing interest declaration

The authors declare no competing interests.

## Notes

### Competing Interest Statement

The authors have declared no competing interest.

