## Supplementary Figures 1-10 for "Mitochondria decode firing frequency and coincidences of postsynaptic APs and EPSPs"

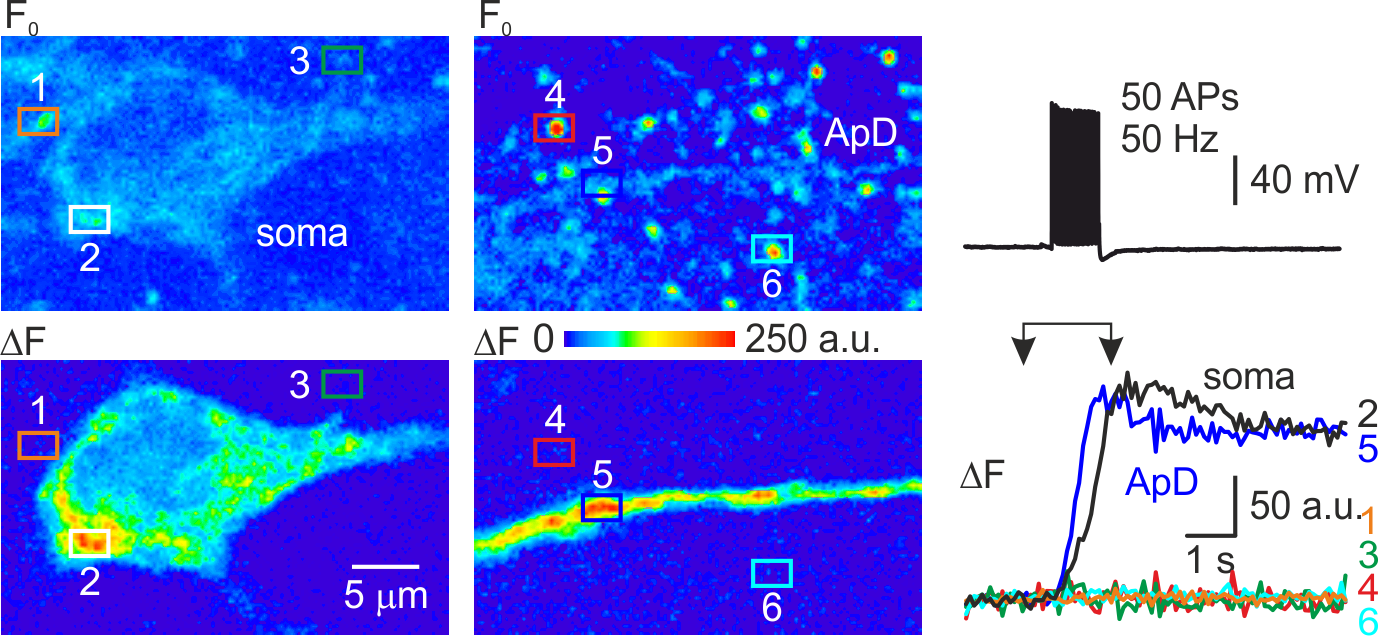
**Supplementary Figures**

**Supplementary Figure 1.** Ca^2+^ elevations were observed only in the mitochondria of the electrically active neurons.

*Left*, Pseudocolor maps of the mitoGCaMP6m resting fluorescence (Fo, top) and of the change in fluorescence elicited by a train of 50 APs at 50 Hz (ΔF, bottom) between the times marked by the arrowheads in the right panel. The rectangles 1-6 indicate the regions of interest from which fluorescence measurements were obtained. ROIs 2 and 5 contain a single mitochondrion within the soma and apical dendrite, respectively, of the electrically active, patched neuron. ROIs 1,3,4,6 contain mitochondria belonging to the nearby, electrically inactive cells.  *Right*, Action potentials elicited by a train of current pulses injected via the somatic whole-cell pipette only trigger Ca^2+^ elevation in the mitochondria of the electrically active neuron (ROI 2, 5).

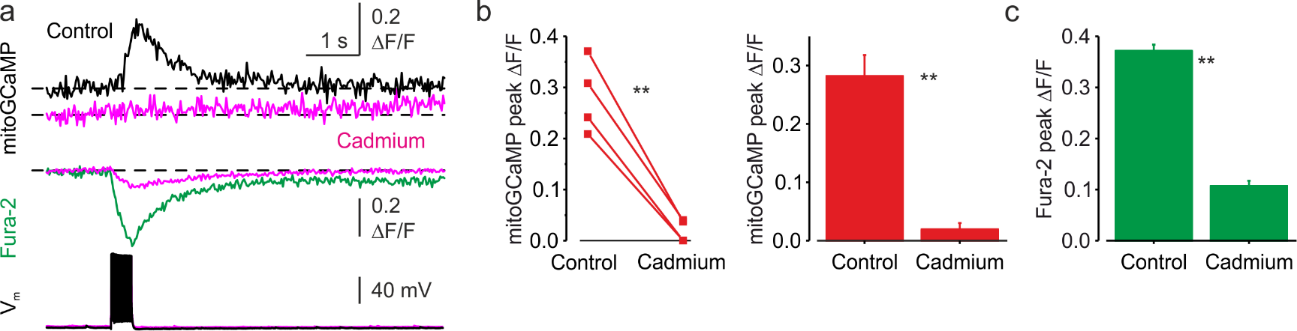

**Supplementary Figure 2.**  Blockade of voltage-gated Ca^2+^ channels abolishes spike-evoked mitochondrial Ca^2+^ transients.

**(a)** Somatic mitoGCaMP6m (*top*) and Fura-2 (*bottom*) transients elicited by a train of 20APs before (black and green respectively) and, 20 minutes after (magenta) bath application of Cadmium (200 µM).

**(b)** Cadmium application blocks mitochondrial Ca^2+^ transients.  *Left,* Peak amplitude mitoGCaMP6m ΔF/F transients elicited by 20 AP before and after application of Cadmium. Notice that the mitochondrial Ca^2+^ transients were almost completely abolished when Cadmium is present. A line connects the paired values obtained from the same individual neuron. Right, Mean peak amplitude of mitoGCaMP6m ΔF/F transients in control*,* after the Cd^2+^ application (28±4% vs. 2±1%, mean±SE, n=4, p<0.01).

**(c)** Cadmium application blocks cytosolic Ca^2+^ transients. Mean peak amplitude of Fura-2 ΔF/F transients in control*,* and after the Cd^2+^ application was significantly suppressed (37±1% vs. 11±1%, mean±SE, n=3, p<0.01).

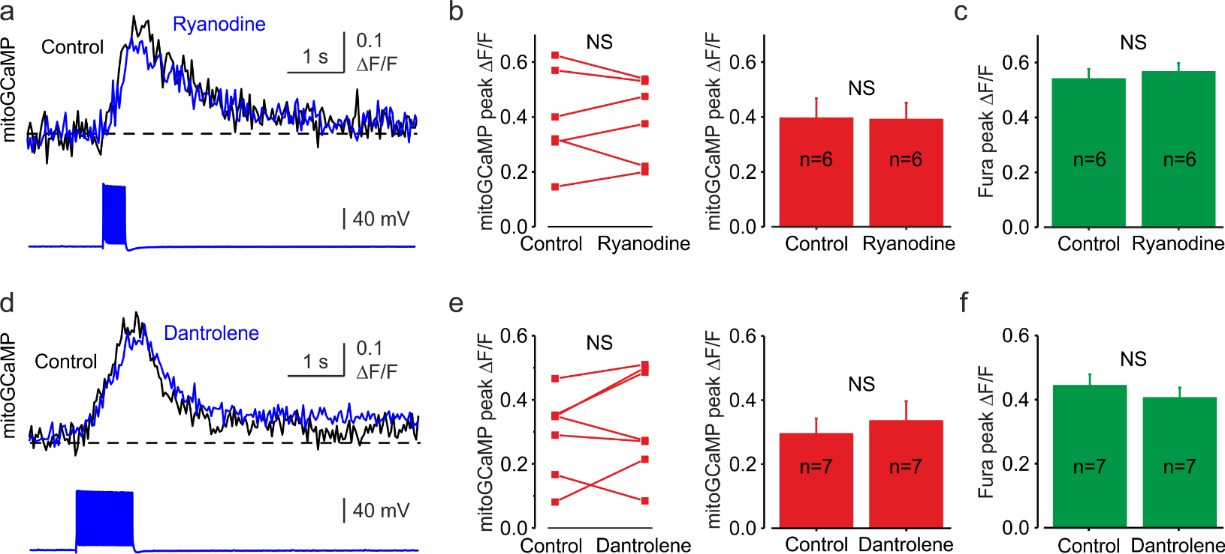

**Supplementary Figure 3.** Blockade of the ER Ryanodine receptors has no significant effect on mitochondrial Ca^2+^ transients.

**(a)** Somatic mitoGCaMP6m transients elicited by a train of 20APs before (black) and 20 minutes after (blue) bath application of high concentration of Ryanodine (100 µM).

**(b)** Blockade of Ryanodine receptor-mediated calcium release from the endoplasmic reticulum did not affect mitochondrial Ca^2+^ transients.  *Left,* The peak amplitude of mitoGCaMP6m ΔF/F transients before and after application of Ryanodine. A line connects the paired values obtained from the same individual neuron.  *Right*, Mean peak amplitude of mitoGCaMP6m ΔF/F transients in control (39±7%, n=6) and after the Ryanodine application (39±6%, n=6, p=0.88)

**(c)** Mean peak amplitude of Fura-2 ΔF/F transients in control and after the Ryanodine application (53±4% vs. 57±3%, respectively, n=6, p=0.37). Notice that Ryanodine produces no significant change in the amplitude of the cytosolic Ca^2+^ transients.

**(d)** Somatic mitoGCaMP6m transients elicited by a train of 50APs before (black) and 20 minutes after (blue) bath application of Ryanodine receptor blocker, Dantrolene (100 µM).

**(e)** Blockade of Ryanodine receptor-mediated calcium release from the endoplasmic reticulum by Dantrolene did not affect mitochondrial Ca^2+^ transients.  *Left,* The peak amplitude of mitoGCaMP6m ΔF/F transients before and after application of Dantrolene. A line connects the paired values obtained from the same individual neuron.  *Right*, Mean peak amplitude of mitoGCaMP6m ΔF/F transients in control (29±5%, n=7) and after the Dantrolene application (33±6%, n=7, p=0.33)

**(f)** Mean peak amplitude of Fura-2 ΔF/F transients in control and after the Dantrolene application (44±4% vs. 40±3%, respectively, n=7, p=0.16). Notice that Dantrolene produces no significant change in the amplitude of the cytosolic Ca^2+^ transients.

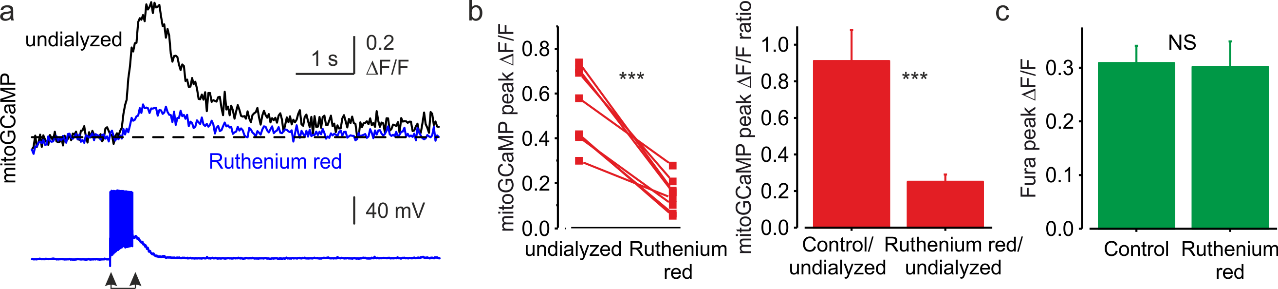

**Supplementary Figure 4**. Intracellular application of MCU blocker, Ruthenium red, abolishes mitochondrial Ca^2+^ transients.

**(a)** MitoGCaMP6m Δ*F*/*F* transients elicited by a train of 20 suprathreshold synaptic stimuli immediately before (black) and 15 minutes after break-in (blue) to the whole-cell configuration with a pipette filled with MCU blocker, Ruthenium red (10 µM). Arrows indicate the beginning and end of the train of stimuli.

**(b)**  The peak amplitudes of the mitochondrial Ca^2+^ transients elicited by a train of 20 suprathreshold synaptic stimuli.  *Left,* the peak amplitude of the dendritic mitoGCaMP6m ΔF/F transients elicited by 20 synaptic stimuli immediately before and 15 minutes after the break-in with Ruthenium red-containing pipette. Notice the mitochondrial Ca^2+^ transient was almost completely abolished after applying Ruthenium red (ΔF/F of 51±1% vs. 13±2%, n=10, p<0.001). A line connects the paired values obtained from the same individual neuron. *Right,* Mean ratio of peak amplitudes mitoGCaMP6m ΔF/F transients obtained from undialyzed neuron and the same neuron after 15 minutes of dialysis with either Ruthenium red containing or control intracellular solution. Dialysis with control solution produced no significant change in the amplitude of the mitoGCaMP6m ΔF/F transients (the ratio of 91±17%, n=5). In contrast, dialysis of Ruthenium red reduced the peak amplitude significantly (the ratio of 25±4%, n=10).

**(c)** Mean peak amplitude of Fura-2 ΔF/F transients 15 minutes after intracellular dialysis with either control or Ruthenium red containing solution. Notice that the cytosolic Ca^2+^ transients are not significantly affected by Ruthenium red dialysis (peak ΔF/F 31±3%, n=14 in control vs. 30±5% in Ruthenium red recordings, n=8, p=0.90).

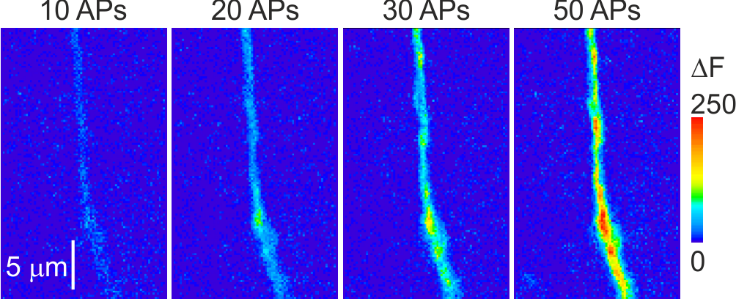

**Supplementary Figure 5.** The amplitude of mitoGCaMP6m transients varies as a function of the number of APs. Shown are pseudocolor maps of change in the mitoGCaMP6m fluorescence in response to trains of 10, 20, 30, and 50 APs at 50 Hz. Fluorescence was measured in the same apical dendrite as in **Supplementary Figure 1**. Notice that the amplitude of the ΔF transients increases as a function of the number of spikes in the train.

**Supplementary Figure 6.** *Left*, double exponential time course of mitoGCaMP6m transient elicited by 50 APs during a prolonged (35 s) optical recording.
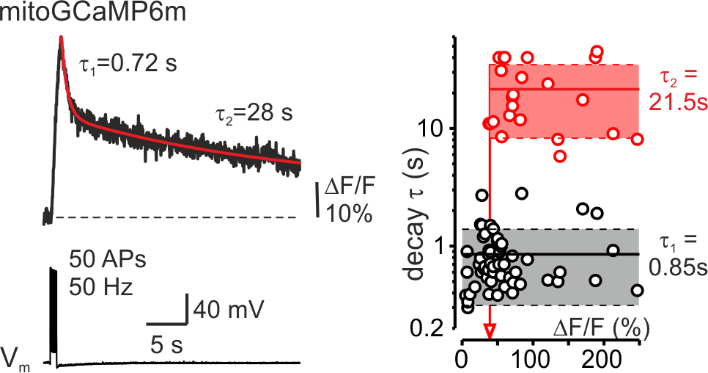
 Right, a plot of the relationship between the decay time constants and the peak amplitude of the mitoGCaMP6m transient. Each dot represents a decay time constant (τ_1_ for monoexponential decays, τ_1_ and τ_2_ for bi-exponential decays) obtained in recordings from 42 neurons. Continuous lines are mean τ_1_ (n= 55, black) and τ_2_ (n=23, red); dashed lines represent standard deviation from the mean. Arrow represents the amplitude of the smallest mitoGCaMP6m transient, which decayed biexponentially.

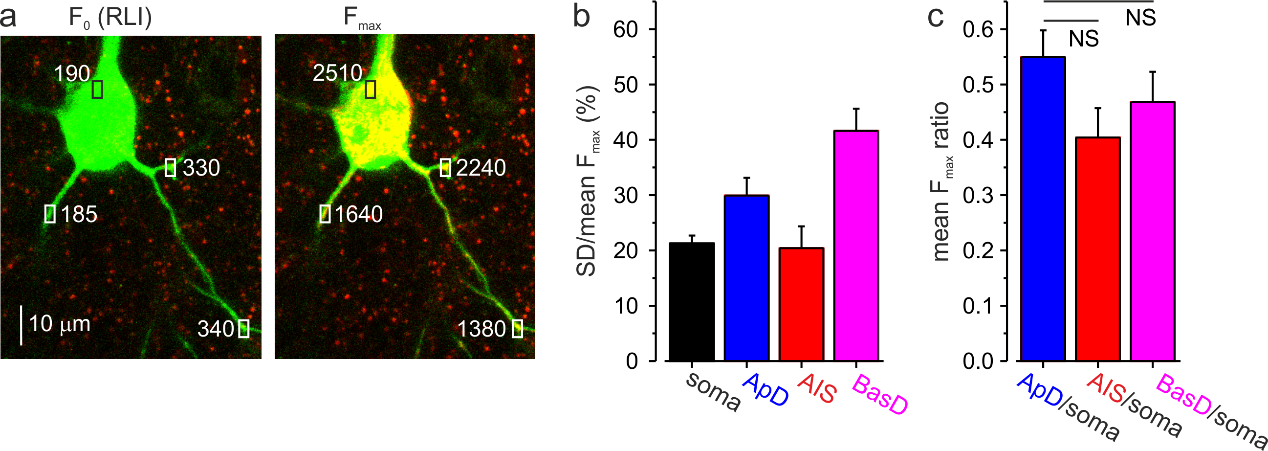

**Supplementary Figure 7.**  Analysis of the resting and maximal mitoGCaMP6m fluorescence reveals its homogenous expression in soma and processes of L5 pyramidal neurons.

**(a)** Individual mitoGCaMP6m labeled mitochondria in a representative voltage-clamped L5 pyramidal neuron at a holding potential of -70 mV and following a one-minute long voltage step +20 mV. The images were obtained by merging the single, 1 µm thick optical sections at excitation wavelengths of 760 and 960 nm, eliciting the Fura-2 (green) and mitoGCaMP6m (red) fluorescence, respectively. An increase in yellow color intensity following the prolonged voltage step reflects an increased mitoGCaMP6m fluorescence. The rectangles indicate the regions within the thin basal dendrite and somata, from which fluorescence measurements were obtained. Notice that the prolonged depolarization elicited a 4-12 times increase in mitoGCaMP6m fluorescence in all neuronal compartments. The smaller spike produced Ca^2+^ elevations in the mitochondria localized in the thin neuronal processes reflect reduced Ca^2+^ influx via the MCU rather than reduced mitoGCaMP6m presence.

**(b)** Variance analysis of the maximal mitoGCaMP6m fluorescence (F_max_) reveals little difference in mitoGCaMP6m expression in the individual mitochondria in the soma and neuronal processes. F_max_ values were measured from 5-70 individual mitochondria in soma, apical and basal dendrites and the AIS of 14 neurons following prolonged depolarizing voltage step. The mean SD/mean of F_max_ was 21±1% for somatic mitochondria, 30±3% in apical dendrites, 20±4% in AIS, and 42±4% in basal dendrites. The relatively narrow F_max_ distributions reflect homogeneity in mitoGCaMP6m expression within the mitochondrial population.

**(c)** Mean ratio of F_max_ in the neuronal process vs. soma. The mean F_max_ ratio was 0.55±0.05 for mitochondria in apical dendrites vs. soma, 0.40±0.05 for mitochondria in AIS vs. soma 0.47±0.05 for mitochondria in basal dendrites vs. soma (mean±SE, n=14 neurons). The smaller F_max_ in neuronal processes might be explained by the smaller depth of the individual mitochondrial rather than by lower mitoGCaMP6m expression.

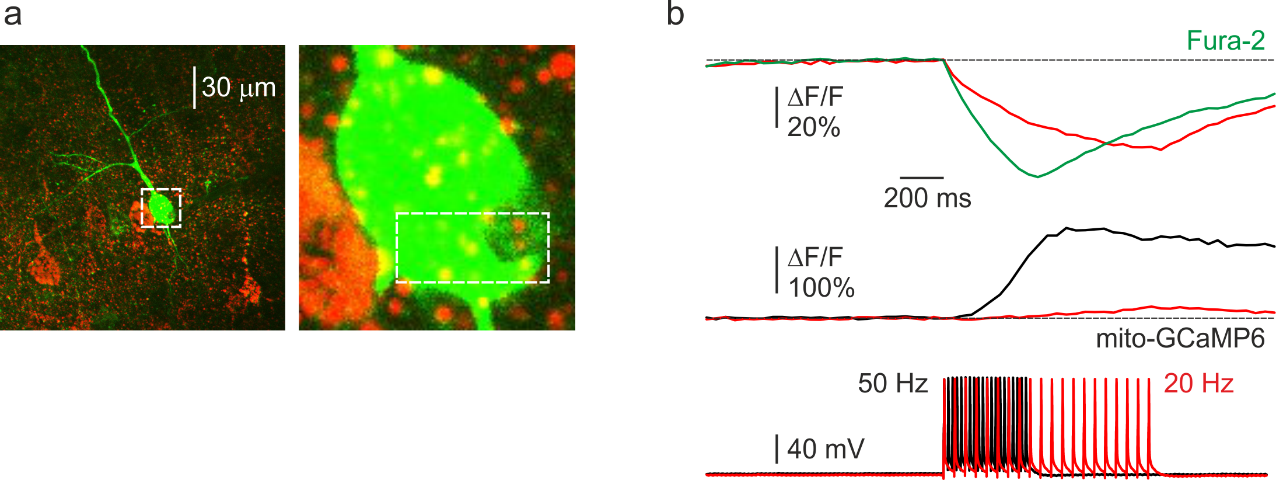

**Supplementary Figure 8.**  The amplitude of the mitochondrial Ca^2+^ elevation elicited by spike train is proportional to the rate-of-rise of the cytosolic Ca^2+^ concentration.

**(a)**  *Left,* an image of a representative L5 pyramidal neuron, obtained by merging the Fura-2 and mitoGCaMP6m fluorescence (for detail, see Figure 1). *Right,* inset: higher magnification image, corresponding to the rectangle in the left panel. The rectangle indicates the region from which the fluorescence measurements were obtained.

**(b)** Δ*F*/*F* transients elicited by a train of 20AP at 20 Hz (red) and 50 Hz (black and green). Cytosolic (top) and mitochondrial (bottom) Ca^2+^ transients. Notice the slower time course of the cytosolic calcium transient in response to a 20 Hz spike train. The slower rise, and not the difference in [Ca^2+^]_i_ level, correlates with the mitochondrial calcium transient amplitude.

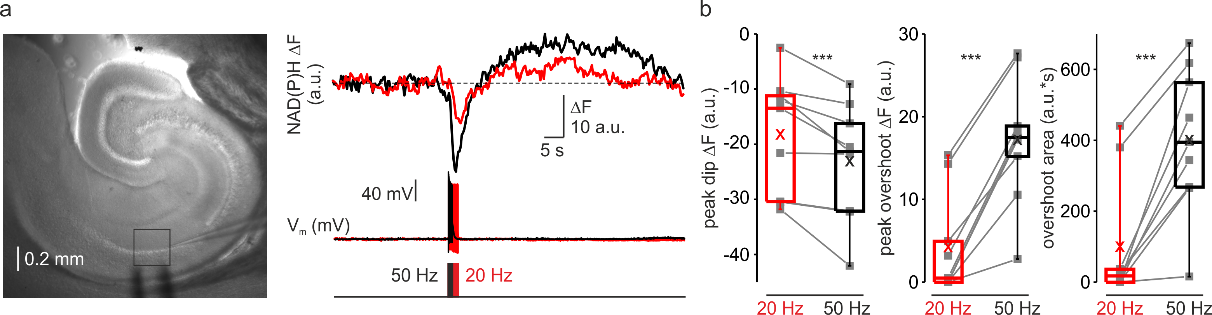

**Supplementary Figure 9.** Frequency-dependent amplification of spike elicited changes in mitochondrial NAD(P)H autofluorescence in the CA1 area of hippocampus.

**(a)** In a representative hippocampal slice, changes in NAD(P)H fluorescence in response to extracellular stimuli trains depend on stimulation frequency.  Left, DIC image of a hippocampal slice during the electrical and optical recording. The rectangle indicates the region from which the auto-fluorescence measurements were obtained. The stimuli were delivered via the bipolar electrode placed in Stratum oriens, and the whole-cell recording was obtained from a nearby CA1 pyramidal neuron. Right, The membrane potential and optical traces evoked by trains of 50 just suprathreshold stimuli at 50 Hz (black) and 20 Hz (red). Notice that both dip and overshoot of the NAD(P)H signals are more prominent at 50 Hz.

**(b)** Higher frequency stimulation causes an increase in the magnitude of the dip and of the overshoot of the NAD(P)H signal. Box plots representing the peaks of the dip (left, -18±4 and -23±4 a.u. for 20 and 50 Hz, respectively, n=9, p<0.005), the peaks of the overshoot (middle, 4±2 and 17±3 a.u. for 20 and 50 Hz, respectively, n=9, p<0.001), and the area of the overshoot (right, 100±59 and 401±69 a.u.*s for 20 and 50 Hz, respectively, n=9, p<0.001) of the NAD(P)H signals. The grey lines connect the paired values obtained from the same regions at two firing frequencies. Box plots represent the 25–75% interquartile range, and the whiskers expand to the 5–95% range. A horizontal line inside the box represents the median of the distribution, and the mean is represented by a cross symbol (X).

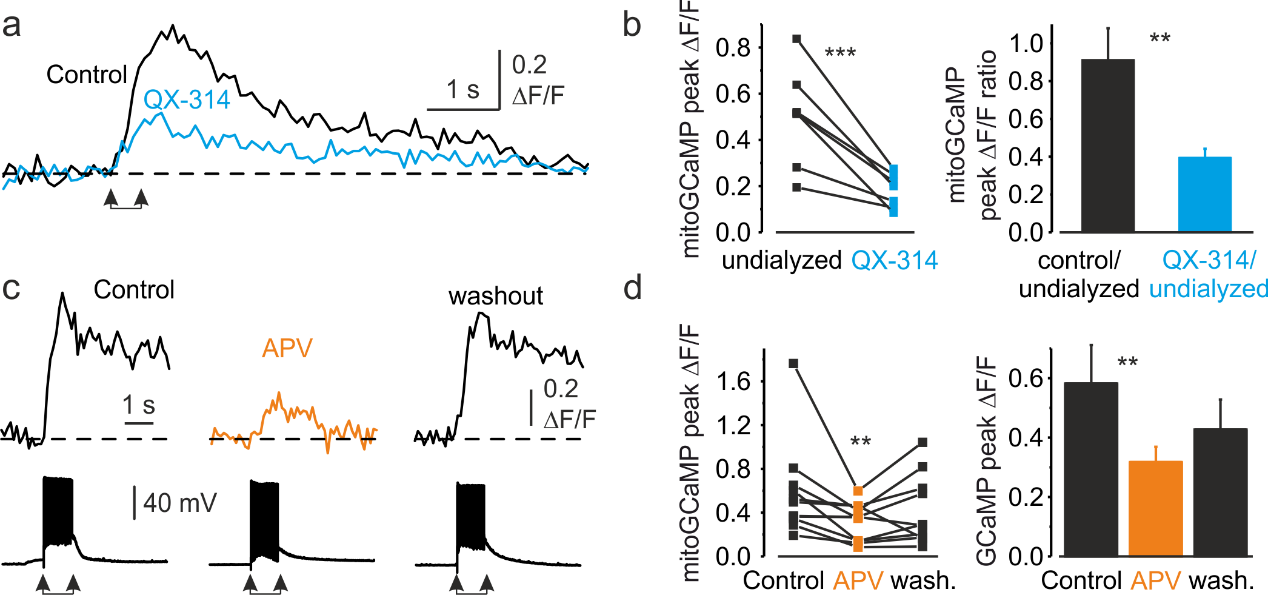

**Supplementary Figure 10.** Availability of postsynaptic Na^+^ channels and synaptic NMDA receptors is required for induction of dendritic mitochondrial Ca^2+^ elevation.

**(a)** MitoGCaMP6m Δ*F*/*F* transients in the apical dendrite elicited by a train of 20 suprathreshold synaptic stimuli before (black) and 10 minutes after a (blue) break-in to the whole-cell configuration with a pipette filled with Na^+^ channel blocker, QX-314 (100 µM). Arrows indicate the beginning and end of the train of stimuli.

**(b)**  The amplitude of the mitochondrial Ca^2+^ elevation elicited in dendrites by trains of synaptic stimuli decreases when QX-314 blocks postsynaptic AP generation.  *Left,* The peak amplitude of the dendritic mitoGCaMP6m ΔF/F transients elicited by 20 synaptic stimuli before (black) and after (blue) intracellular application of QX-314. Notice a significant decrease in amplitude of mitochondrial Ca^2+^ signals when AP generation is blocked (from 50±8% to 18±3%, n=7, p<0.005). A line connects the paired values obtained from the same individual neuron. *Right,* Mean ratio of peak amplitudes of synaptically elicited, dendritic mitoGCaMP6m ΔF/F transients after the dialysis with either control (black, n=8) or QX-314-containing intracellular solution (blue, n=7) to those before the establishment of whole-cell recording. Notice that while the dialysis with control solution elicits little change in the amplitude of mitochondrial Ca^2+^ signals, blockade of postsynaptic AP causes a significant decrease in the ratio (p<0.005).

**(c)** MitoGCaMP6m Δ*F*/*F* transients in a representative apical dendrite before, after bath application of APV (50 µM), and following the washout. The bottom traces represent somatic membrane potential. Arrows mark the beginning and end of the synaptic stimuli train.

(**d)** Blockade of NMDA receptors causes a decrease in amplitude of the synaptically evoked mitochondrial Ca^2+^ transients in dendrites.  *Left,* The peak amplitude of the dendritic mitoGCaMP6m ΔF/F transients elicited by 20 synaptic stimuli before, during APV application, and upon the washout. Notice a significant decrease in the transients' amplitude (n=10, p<0.01) when APV is present. A line connects the paired values obtained from the same individual neuron. *Right*, Mean peak amplitude of mitoGCaMP6m ΔF/F transients in control*,* after the APV application and after the washout (58±13%, 32±5% and 43±10%, respectively, n=10).
